# Phylogenetic origin and sequence features of MreB from the wall-less swimming bacteria *Spiroplasma*

**DOI:** 10.1101/2020.05.13.095042

**Authors:** Daichi Takahashi, Ikuko Fujiwara, Makoto Miyata

**Author notes:** Address correspondence to Makoto Miyata.

## Abstract

*Spiroplasma* are wall-less bacteria which belong to the phylum Tenericutes that evolved from Firmicutes including *Bacillus subtilis. Spiroplasma* swim by a mechanism unrelated to widespread bacterial motilities, such as flagellar motility, and caused by helicity switching with kinks traveling along the helical cell body. The swimming force is likely generated by five classes of bacterial actin homolog MreBs (SMreBs 1-5) involved in the helical bone structure. We analyzed sequences of SMreBs to clarify their phylogeny and sequence features. The maximum likelihood method based on around 5,000 MreB sequences showed that the phylogenetic tree was divided into several radiations. SMreBs formed a clade adjacent to the radiation of MreBH, an MreB isoform of Firmicutes. Sequence comparisons of SMreBs and *Bacillus* MreBs were also performed to clarify the features of SMreB. Catalytic glutamic acid and threonine were substituted to aspartic acid and lysine, respectively, in SMreB3. In SMreBs 2 and 4, amino acids involved in inter- and intra-protofilament interactions were significantly different from those in *Bacillus* MreBs. A membrane-binding region was not identified in most SMreBs 1 and 4 unlike many walled-bacterial MreBs. SMreB5 had a significantly longer C-terminal region than the other MreBs, which possibly forms a protein-protein interaction. These features may support the functions responsible for the unique mechanism of *Spiroplasma* swimming.

## 1. Introduction

The bacterial phylum Tenericutes that evolved from the phylum Firmicutes is comprised of the class Mollicutes, which includes the genera *Spiroplasma* and *Mycoplasma* [1]. Species of these genera are characterized by small genome size, being mostly parasitic or commensal, and absence of peptidoglycan. In addition, they cannot equip conventional machineries for bacterial motility, such as flagella or type IV pili, due to the lack of a peptidoglycan layer. Instead, they have developed three unique motility systems: *mobile* type gliding, *pneumoniae* type gliding, and *Spiroplasma* swimming [2].

*Spiroplasma* have a helical cell shape and infect plants and arthropods [3,4]. They swim by rotating the cell body in viscous environments, including host tissues. To rotate the cell body, *Spiroplasma* change the helicity handedness at the front end by forming a kink, and propagate the kink along the cell body to back (Fig. 1A&B) [5,6,7,8]. The helical cell shape originates from a flat ribbon structure (Fig. 1C) extending along the innermost line of the cell, and kink propagation is likely caused by a structural change of the ribbon [5,9,10,11]. Electron microscopy and proteome analyses have revealed that the flat ribbon is composed of fibril, a *Spiroplasma* specific protein that forms filaments, and MreB proteins, which are bacterial actin homologs (Fig. 2A) [9,10,11]. Tenericutes lack respiration pathway to generate membrane potential, and produce ATP through glycolysis and arginine fermentation [12]. Then, the energy for cell activities is mostly supplied from ATP rather than membrane potential in many species. Actually, the energy for *mobile* and *pneumoniae* type gliding mechanisms is supplied from ATP [2]. *Spiroplasma* swimming may also be based on ATP energy. Regarding the component proteins of the ribbon-shaped swimming machinery, fibril is related to adenosyl homocysteine nucleosidase that cannot hydrolyze ATP [13]. On the other hand, MreB is well known as an ATPase like other actin superfamily proteins [14,15]. These facts suggest that the MreBs are employed in the *Spiroplasma* swimming.

**Fig. 1.**
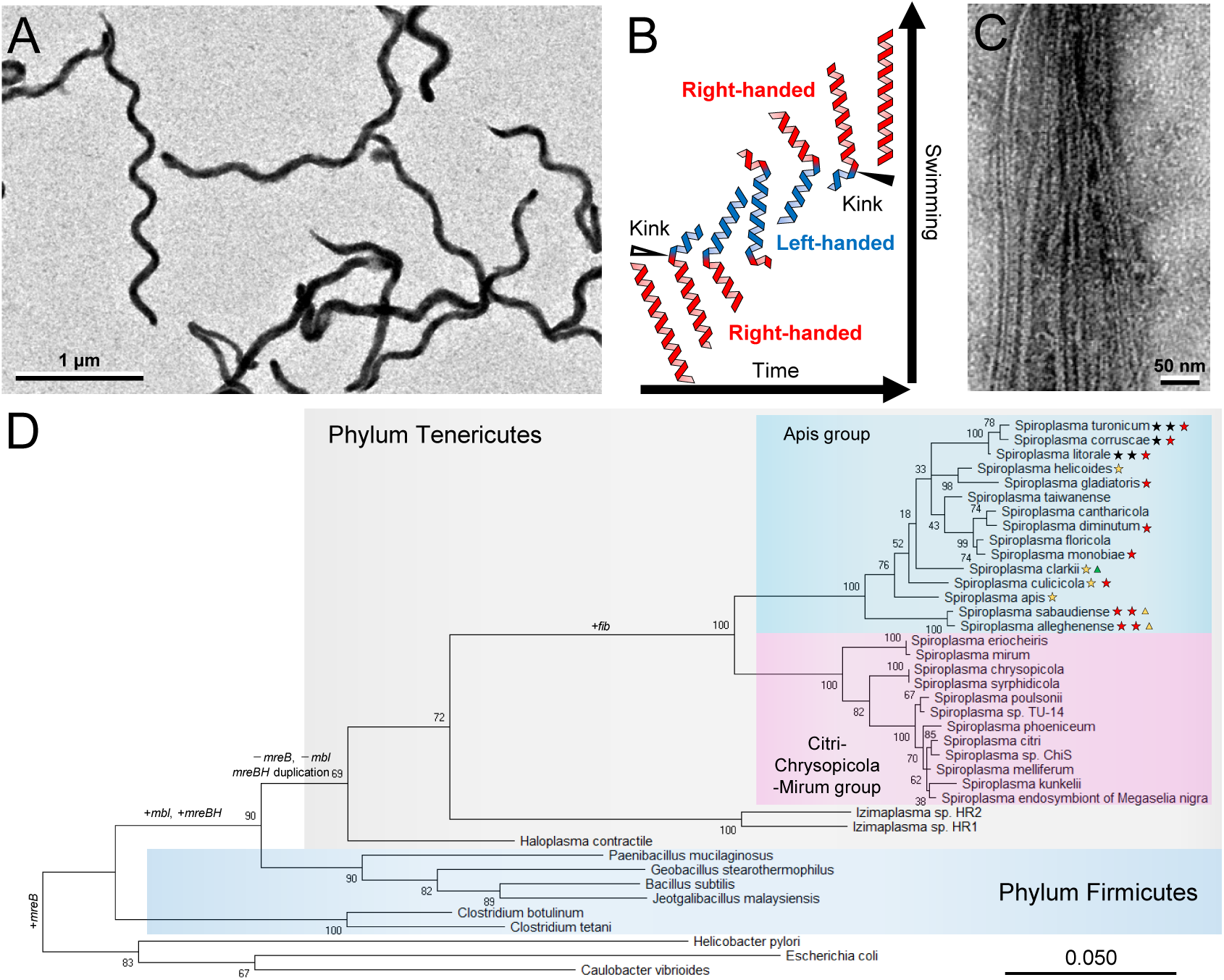
Morphologies, swimming and phylogeny of *Spiroplasma*. (A) Electron microscopy of *Spiroplasma eriocheiris*. (B) Swimming scheme. *Spiroplasma* swim via the rotation of the cell body caused by kink propagation along the cell axis from front to back, which causes helicity shifts of right to left (open triangle) and left to right (closed triangle). (C) Electron microscopy of filaments from ribbon. (D) Maximum likelihood phylogeny for 16S rRNA of the organisms that are related to the evolution of *Spiroplasma*. Sequences for 16S rRNA of proteobacteria (*E. coli, Caulobacter vibrioides*, and *Helicobacter pylori*) were used as the outgroup. Bootstrap support values are indicated on each node. Stars and triangles marked for each species indicate one duplication and deletion, respectively for genes of SMreBs colored differently as yellow: SMreB1, green: SMreB2, black: SMreB4, and red: SMreB5. The scale bar is in units of the number of nucleotide substitutions per site. Panels (A) and (C) were modified from a previous paper [9].

**Fig. 2.**
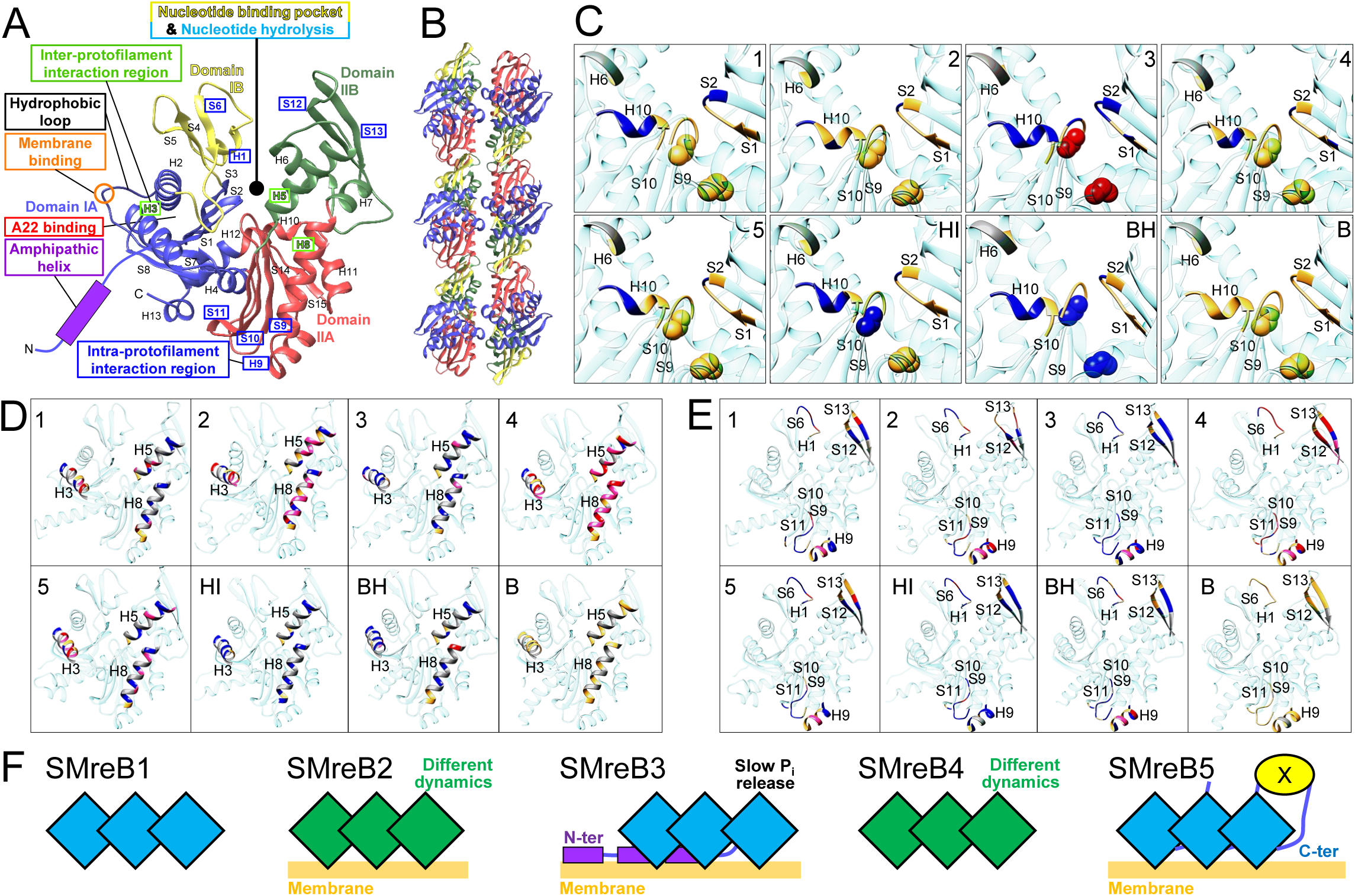
Structural features of MreB proteins. (A) Structure of *Bacillus subtilis* MreB predicted from the amino acid sequence (NP_390681.2). The structure is composed of 15 strands and 13 helices shown as S1-S15 and H1-H13, respectively. Individual domains are colored blue (IA), yellow (IB), red (IIA), and green (IIB). An N-terminal amphipathic helix carried by some Gram-negative bacterial MreBs is presented as a rectangle colored in purple. The nucleotide binding pocket, catalytic region for nucleotide hydrolysis, A22 binding region, inter- and intra-protofilament interaction regions, hydrophobic loop, membrane binding amino acids in the hydrophobic loop, and N-terminal amphipathic helix are marked with a rectangle common in style to those in Fig. 4. (B) Filamentous structure of MreB extracted from three consecutive unit cells of a *Caulobacter crescentus* MreB crystal (PDB ID: 4CZJ). (C-E) Focused amino acids are shown as 3D structures. The structures for SMreBs 1-5, HMreB, BMreBH, and BMreB were marked as 1-5, HI, BH, and B, respectively. The amino acids were colored red, pink, blue, or gold, based on the conservation levels compared to BMreBs (red: conserved amino acids whose corresponding ones in BMreB were conserved as different amino acids, pink: conserved amino acids whose corresponding ones in BMreB were not conserved, blue: not conserved amino acids whose corresponding ones in BMreB were conserved, gold: conserved amino acids whose corresponding ones in BMreB were conserved as the same amino acid). The numbers of helices and strands are indicated for colored parts. (C) Nucleotide binding pocket. The backbone atoms of catalytic amino acids for nucleotide hydrolysis are indicated by spheres. (D) Inter-protofilament interaction region. (E) Intra-protofilament interaction region. (F) Summary of predicted functions, presenting SMreB molecules (squares), slow P_i_ release (only statement), different polymerization dynamics (green color of squares), membrane binding (yellow thick line), N-terminal membrane binding region (purple rectangle), C-terminal interacting region (blue line), and a putative interacting protein (yellow ellipse). Membrane binding and N-terminal membrane binding region are presented based on Citri group SMreBs, and the others are presented based on both Apis and Citri groups.

Generally, as seen in eukaryotic actin, the MreB molecule has a U-shaped structure that can be divided into four domains (Fig. 2A) [15]. The groove of the U-shaped structure functions as a nucleotide-binding pocket, such as for ATP and GTP (Fig. 2A, S3A) [15]. A previous study reported that MreB polymerizes into non-helical filaments with juxtaposed subunits in which two protofilaments interact antiparallelly in the presence of nucleotides (Fig. 2B) [15]. Molecular dynamics (MD) simulations suggest that the inter- and intra-protofilament interactions are comprised of four helices, two strands, and three loops (H3, H5, and H8 shown in Fig. 2A and Fig. S3C for inter-protofilament interaction; H9, S12, S13 and loops between S6-H1, S9-S10, and S10-S11 shown in Fig. 2A and Fig. S3D for intra-protofilament interaction) [16]. Unlike the eukaryotic actin filament, MreB filaments bind to the cell membrane directly [17]. A previous study reported that many MreBs have two consecutive hydrophobic amino acids for membrane binding in the hydrophobic loops in domain IA, and some MreBs of Gram-negative bacteria have an amphipathic helix at the N-terminus as an additional factor for membrane binding (Fig. 2A) [17].

In various phyla of rod-shaped bacteria, MreB is well-conserved in the genomes and is responsible for cell shape maintenance [18]. The MreB filament binds to the membrane to form an elongasome complex with other proteins, which precisely locates the peptidoglycan-synthesizing enzymes to retain the rod shape of the cells [18]. Many bacterial species represented by *Escherichia coli* have a single copy of the *mreB* gene on their genome [18], whereas the genome of *Bacillus subtilis* has three genes to code MreB isoforms (MreB, Mbl, and MreBH), all of which are employed in cell shape maintenance [19].

As *Spiroplasma* MreBs (SMreBs) may have a role that is distinct from those of other MreBs, sequence analyses among SMreBs and walled-bacterial MreBs are useful for clarifying the origin and mechanism of *Spiroplasma* swimming. Moreover, amino acid sequences specific for MreB functions have not yet been characterized for SMreBs.

In this study, we analyzed the sequences of SMreBs. Phylogenetic analyses suggested that SMreBs evolved from MreBH in Firmicutes. Sequence comparisons of SMreBs and *Bacillus* MreBs clarified their sequence features for ATP hydrolysis, interactions for filament formation, and membrane binding.

## 2. Materials and methods

### 2.1. Phylogenetical analyses

A protein BLAST search on non-redundant protein sequences was performed to collect 5,000 amino acid sequences of MreB in ascending order of E value (the maximum value was calculated as 2×10^−63^ for MAV90490.1) by templating MreB3 of *S. eriocheiris* (WP_047791951.1) on July 14, 2019. Sequences of *B. subtilis* MreB isoforms annotated to MreB (AQR87043.1) and MreBH (BAA07047.1) were added to the 5,000 sequences as representative sequences. The sequence alignments and construction of phylogenetic trees were performed via the MUSCLE program, and maximum likelihood and Neighbor-joining methods, respectively, using MEGA-X ver. 10.1 [20]. Bootstrap supports were estimated from 100 samples of alignment. For the ancestor estimation, maximum likelihood phylogenetic trees by each MreB group were constructed with a *B. subtilis* MreB (an *E. coli* MreB for the estimation of *Bacillus* MreB (BMreB)) as an outgroup, and the ancestral sequences were estimated by the ancestor estimation program of MEGA-X ver. 10.1 [20].

### 2.2. Structure prediction and sequence analyses

All monomeric MreB structures in this study were predicted using Rosetta comparative modeling by templating a structure of *Caulobacter crescentus* MreB (PDB ID: 4CZJ), using Robetta software [15,21]. Sequence comparisons among amino acid sequences were performed by Clustal Omega [22]. Duplicated SMreBs 1, 4, and 5 in the Apis group species were incorporated into the analyses. Seven of the 19 amino acid sequences contained in SMreB3 of the Apis group had no annotation. Since these proteins were coded in a position suggesting SMreB3 as an ortholog on the genome, they were included in the analyses. SMreB3 of the *S. melliferum* strain CH-1 (AHA83319.1) was excluded from the analyses because it was approximately 40% shorter than the other SMreB3 sequences. Two sequences, MreB4 of *S. eriocheiris* strain CCTCC_M_207170 (AHF58342.1) and a protein from *S. culicicola*, which is unannotated but can be classified to MreB3 (WP_025363622.1), were not used for the analyses because the initiation codons were miss-assigned. The predictions of secondary structures and amphipathic helices were performed using PSIPRED 4.0 and AmphipaSeeK software, respectively [23,24]. Sequence identity and similarity were calculated as the ratio of amino acids with identity and strong similarity defined by the Gonnet PAM 250 matrix over the total amino acid number excluding gap regions.

## 3. Results and discussion

### 3.1. Evolutional relationships between Spiroplasma and other MreBs

We analyzed the phylogenetic relationships among SMreBs and other MreBs. We first obtained 5,002 amino acid sequences of MreB family proteins through a BLAST search, including all 170 non-redundant sequences of SMreB with those of three new species (*S*. sp. TU-14, *S*. sp. ChiS, and *S*. endosymbiont of *Megaselia nigra*) (Table S1) (Data set 1) [3,4,25]. Based on these sequences, we constructed a maximum likelihood phylogenetic tree (Fig. 3, S7). We adopted this phylogenetic tree because the significant difference was not found among the topologies of a maximum likelihood phylogeny using 4,000 sequences, the Neighbor-Joining phylogeny using 5,002 sequences, and the phylogeny shown in Fig. 3 (and Fig. S7). The largest radiation of the phylogenetic tree consisted of MreB family proteins of conventional bacteria, which were further divided into Gram-negative and positive bacteria. Previous studies of MreB family proteins have primarily focused on proteins in these radiations [18]. In our phylogenetic tree, an MreB radiation of candidate phyla radiation (CPR) [1], which is composed of Parcubacteria and Microgenomates superphyla, was formed next to the radiation of conventional MreBs. The CPR MreBs were split into two clades. Three candidate species of Parcubacteria in candidate phyla Yanofskybacteria, Taylorbacteria, and Zambryskibacteria, and a candidate species of Microgenomates in candidate phylum Woesebacteria had MreBs of both clades, suggesting the existence of two types of MreBs possessing different roles in CPR.

**Fig. 3.**
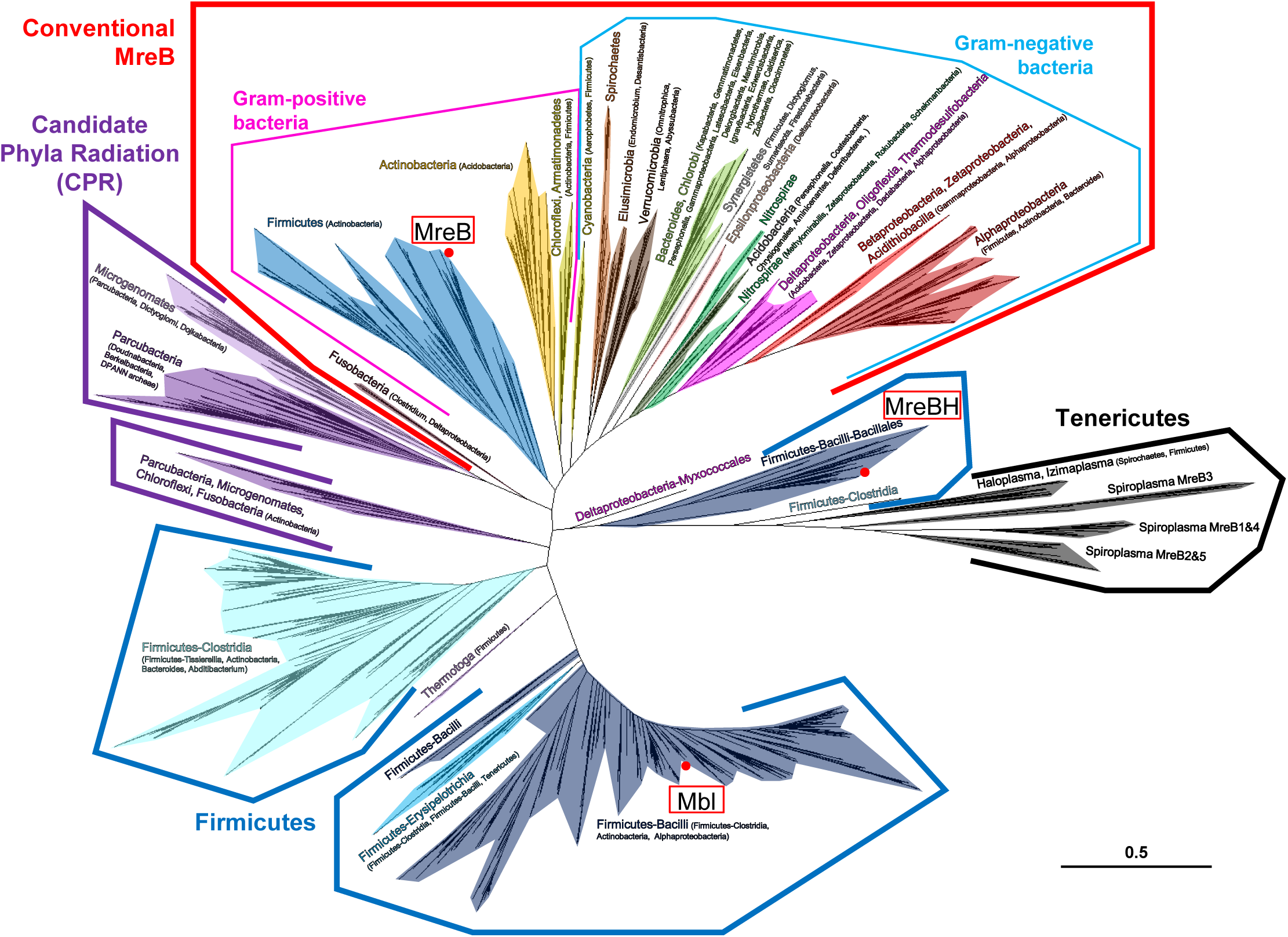
Unrooted maximum likelihood phylogenic tree for 5,002 amino acid sequences of MreB family proteins. The radiation of conventional MreBs, which are conserved over various walled bacteria, is indicated by a red polygon. MreB isoforms of Firmicutes, and MreBs of CPR and Tenericutes form radiations other than conventional MreBs, and are indicated by blue, purple, and black polygons, respectively. Each clade is colored according to the major phyla (or classes). Major and minor taxa in each clade are indicated without and with parentheses, respectively. The branches of MreB isoforms in *Bacillus subtilis* are marked by red circles. The scale bar is in units of the number of amino acid substitutions per site.

Firmicutes bacteria have more than one copy of a gene coding MreB isoforms on the genomes [19], and these isoforms formed three independent clades. One was mostly composed of class Bacilli MreB family proteins related to *Bacillus* Mbl. Another clade positioned between clades of *Bacillus* Mbl and CPR MreB was primarily composed of proteins in class Clostridia, distantly related to *Bacillus* MreB isoforms, suggesting the existence of a new group of MreB isoforms. The clade including *Bacillus* MreBH (BMreBH) had a root distinct from other clades. SMreBs formed a radiation near the BMreBH clade. A clade of MreBs from *Haloplasma* and *Candidatus* Izimaplasma, belonging to Tenericutes phylum (HMreB) was identified between clades of SMreB and BMreBH (Fig. 3, S1). *Haloplasma contractile* is known to perform characteristic motility via tentacle-like structures extending from a coccoidal body [26]. HMreB may be involved in this characteristic motility of *Haloplasma* [26,27], while the motility of Izimaplasma has not been reported yet.

### 3.2. Development of Spiroplasma swimming

Based on the phylogeny results, we assessed the evolutional background prior to *Spiroplasma* acquiring swimming motility using a phylogenetic tree of 16S rRNA, including newly obtained *Spiroplasma* species (Fig. 1D, Data set 2). A Firmicutes species, the ancestor of Tenericutes, probably possessed MreB, Mbl, and MreBH because the Firmicutes species closest to Tenericutes had these three MreB isoforms [1]. The ancestral bacterium then likely lost MreB and Mbl and duplicated MreBH before *Haloplasma* and other Tenericutes bacteria separated. After the duplication of MreBH, *Haloplasma* and Izimaplasma likely acquired HMreBs (Fig. 1D, 3). A previous study suggested that *Haloplasma* and *Spiroplasma* acquired their MreBs by independent gene duplications [27]. However, HMreB was located closer to SMreB and MreBH than to the other clades in our phylogenetic tree (Fig. 3, S1). Accordingly, *Spiroplasma* likely acquired SMreBs through the duplication of MreBH (Fig. 1D). Fibril, another component of the ribbon structure of *Spiroplasma* cells, was likely acquired after *Spiroplasma* and Izimaplasma separated, because it is specific for *Spiroplasma* [9,10,13,27].

The *Spiroplasma* genus can be divided into two subgroups: Apis and Citri-Chrysopicola-Mirum (Citri) [3,25,27]. In terms of the *Spiroplasma* genomes, duplications of SMreBs 1, 4, or 5 and deletions of SMreBs 1 or 2 were identified in almost all species of the Apis group unlike the Citri group (Fig. 1D) [27]. *Spiroplasma* swimming has been studied by using Citri species (*S. melliferum, S. citri*, and *S. eriocheiris*), all of which swim via similar kink propagations [5,7,8,9,10,11,13,28]. The duplication and deletion of SMreBs in the Apis group may suggest their diverse swimming formats.

### 3.3. Sequence comparisons among SMreBs and BMreBs

We compared the amino acid sequences of SMreB, HMreB, BMreB, and BMreBH to characterize SMreBs (Fig. 4, S6). First, the conserved amino acids of SMreBs 1-5, HMreB, and BMreBH sequences were compared with those of 48 sequences of BMreBs (Fig. 4, S6). Each SMreB type showed specific conserved sequences different from those of BMreBs. SMreB 1-5 conserved 128, 187, 46, 210, and 121 amino acids in approximately 330 full-length amino acid sequences, respectively. Of these amino acids, 50, 102, 21, 122, and 53 amino acids were specific for each SMreB, respectively, and were not conserved in BMreBs (Table S2, Fig. 4, S6). Amino acids that are specifically conserved among SMreBs may be essential for intrinsic functions of SMreBs. Sequences of Citri group SMreBs were more conserved than those of the Apis group, corresponding to the phylogenetical spread among *Spiroplasma* species represented in the 16S rRNA phylogeny (Table S3, S4, Fig. 1D, S2, S6).

**Fig. 4.**
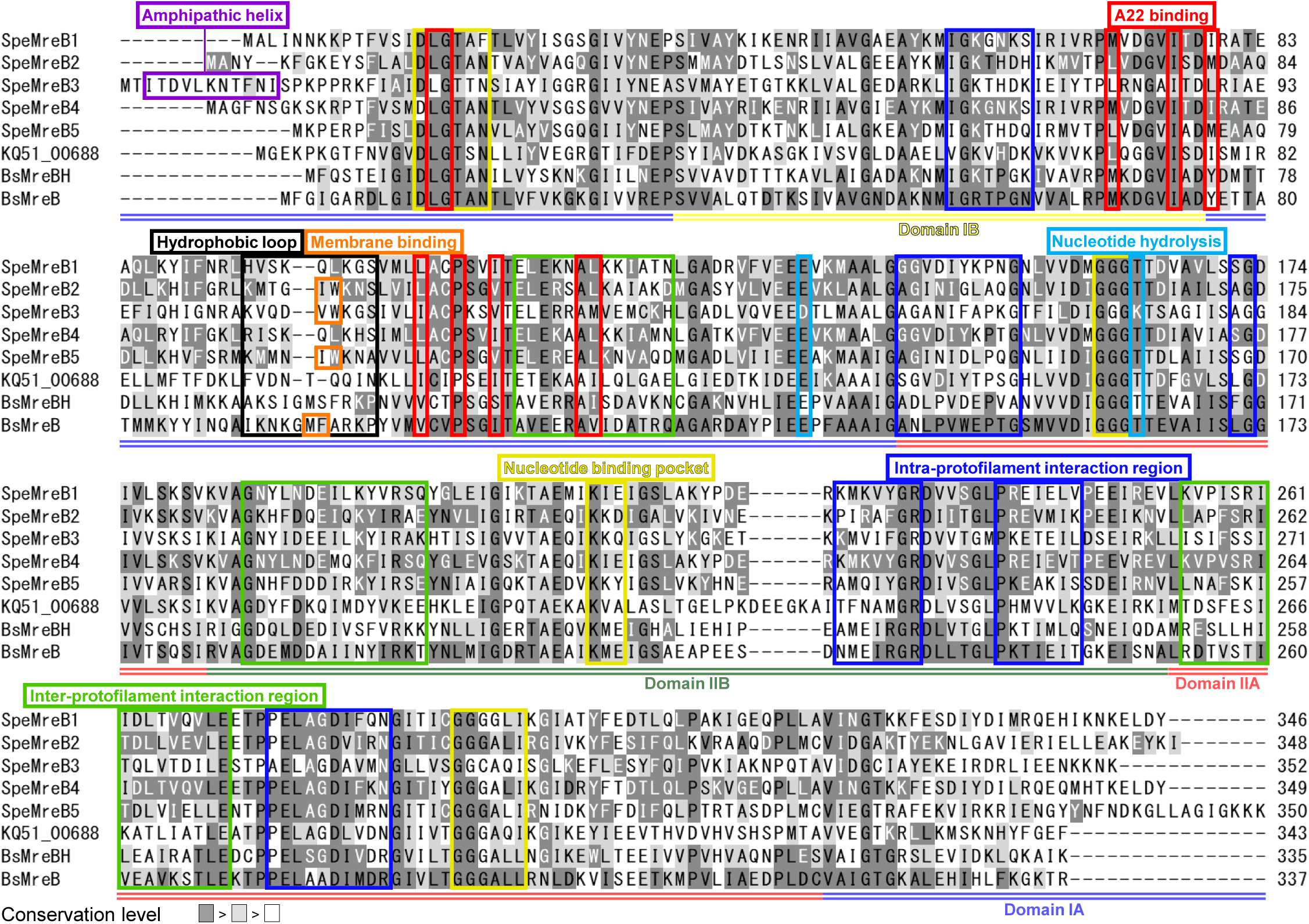
Multiple alignment of SMreBs 1-5, HMreB, BMreB, and BMreBH performed for MreBs 1-5 of *S. eriocheiris*, KQ51_00688 in Izimaplasma sp. HR1, *B. subtilis* MreBH, and *B. subtilis* MreB. The conservation levels for amino acids calculated from multiple alignments of each MreB are indicated by the gradient of gray markers (amino acids calculated as identical and strongly conserved based on the Gonnet PAM 250 matrix are marked by deep and light gray, respectively. The other amino acids are not colored). In SMreBs 1-5, HMreB, and BMreBH, the conserved amino acids, whose corresponding amino acids in BMreB were conserved as a different one or were not conserved, are indicated by white characters. Domains IA, IB, IIA, and IIB are identified by double underlines of coordinated colors with Fig. 2A. Sequences important for the specific function of MreB are indicated by rectangles with different colors.

### 3.4. Characterization of SMreBs based on amino acid sequences

Next, to investigate the SMreB sequences corresponding to the sequences conserved in walled bacteria [15,16,17,29,30], we predicted the ancestral sequences and 3D structures for SMreBs 1-5, HMreB, BMreBH, and BMreB (Fig. 2C-E, S2, Data set 3). First, we focused on the 19 amino acids that construct the nucleotide binding pocket (Fig. 2A&C, S3A) [30]. In SMreB sequences, most of the 11 amino acids conserved in BMreBH were also conserved (Fig. 4 yellow boxes, Fig. 2C, S6). We also focused on the catalytic amino acids for nucleotide hydrolysis [15]. While a glutamic acid and threonine were conserved in almost all MreBs, interestingly, in SMreB3, these sequences were changed to an aspartic acid and lysine, respectively (Fig. 4 cyan boxes, Fig. 2C, S6). As mutations on the catalytic amino acids reduce P_i_ release rates of eukaryotic actin [31], SMreB3 may also have slower P_i_ release rates than other SMreBs (Fig. 2F). It has been reported that an antibiotic A22 inhibits MreB polymerization by interacting with the nucleotide in the MreB molecule [29]. SMreBs also conserved these target amino acids of A22 (Fig. 4 red boxes, Fig. 2A, S3B, S6) [29]. Next, we focused on 87 amino acids involved in the inter- and intra-protofilament interactions (H3, H5, H8, H9, S12, and S13 and loops between S6-H1, S9-S10, and S10-S11 in Fig. 2A, Fig. S3C&D) [16]. In these regions, 31, 50, 14, 69, and 32 amino acids were conserved in SMreBs 1-5, respectively (Fig. 4 green and blue boxes, Fig. 2D&E, S6). In these amino acids, 15, 26, 6, 40, and 16 sequences were not conserved in BMreBs, respectively (white characters in light green and blue boxes in Fig. 4). As these regions in a walled-bacterial MreB move toward an adjacent MreB molecule to interact with each other [16], these amino acid substitutions of SMreBs, especially SMreBs 2 and 4, may affect the flexibility and interaction strength, allowing them to show polymerization characters different to those of walled-bacterial MreBs (Fig. 2F).

We also characterized regions required for membrane binding. Two consecutive hydrophobic amino acids are conserved in many walled-bacterial MreBs (Fig. 2A) [17]. However, SMreBs 1 and 4 of the Citri group did not have hydrophobic amino acids at the corresponding position (Fig. 4 orange boxes, Fig. 2F, S6) [15,17]. Moreover, the consecutive hydrophobic amino acids cannot be found in 46 out of 88 sequences of the Apis group SMreBs 1, 3, 4, and 5 (Fig. S6). These SMreBs may not directly interact with the membrane. We next focused on the N-terminal amphipathic helix found in some MreBs of Gram-negative bacteria (Fig. 2A). All N-terminal regions of the Citri group SMreB3, except for *S. melliferum* strain AS576 (QCO24517.1), were predicted to take amphipathic helices (Fig. 4 purple box, Fig. S4, S6). This suggests that Citri group SMreB3 proteins also have affinity to the membrane through the helix (Fig. 2F).

The lengths of C-terminal sequences varied among 133 SMreBs, and over 79% of SMreBs were eight amino acids longer than that of BMreB (Table S5, Fig. S5, S6). In particular, SMreB5 had the longest C-terminal sequence of all SMreBs (Table S5). Such terminal extension is a feature reminiscent of a bacterial tubulin homolog, FtsZ, featured with a long unstructured region. This is the interaction region for several proteins, including FtsA, ZipA, and ZapD involved in Z-ring formation, which causes cell constriction for bacterial cytokinesis [32]. The extension of the SMreB C-termini may play a role in the protein-protein interactions (Fig. 2F).

## 4. CONCLUSIONS

In this study, we suggested that the five classes of SMreBs evolved from MreBH (Fig. 3) and clarified that the classes can be characterized by ATP hydrolysis, interactions for filament formation, and membrane binding. Based on these results, we suggest the functions for individual SMreBs (Fig. 2F). These would be clue to clarify the origin and mechanism of *Spiroplasma* swimming.

## Conflict of interest

The authors declare no conflict of interests.

## Funding

This study was supported by a Grants-in-Aid for Scientific Research (A and C) (MEXT KAKENHI, Grant Numbers JP17H01544 to MM and JP20K06591 to IF), JST CREST (Grant Number JPMJCR19S5) to MM, Research Foundation of Opto-Science and Technology to IF, and the Osaka City University (OCU) Strategic Research Grant 2019 to IF. DT is a recipient of a Japanese scholarship of JEES Kureha (Toyobo) Scholarship.

## Acknowledgements

We thank Yuya Sasajima and Hana Kiyama at the Graduate School of Science, Osaka City University, Japan for helpful discussion.

## Notes

### Competing Interest Statement

The authors have declared no competing interest.

### Summary of Updates

small modifications for better understanding.

